# Neural Correlates of the Natural Observation of an Emotionally Loaded Video

**DOI:** 10.1101/152298

**Authors:** Melanni Nanni, Joel Martínez-Soto, Leopoldo Gonzalez-Santos, Fernando A. Barrios

**Affiliations:** Universidad Nacional Autónoma de México, Instituto de Neurobiología, Querétaro, México.; Department of Psychology, Universidad de Guanajuato, León, GTO, México.

**Keywords:** stress, brain connectivity, fMRI, resting state networks, free observation, stressful video

## Abstract

Studies based on a paradigm of free or natural viewing have revealed characteristics that allow us to know how the brain processes stimuli within a natural environment. This method has been little used to study brain function. With a connectivity approach, we examine the processing of emotions using an exploratory method to analyze functional magnetic resonance imaging (fMRI) data. This research describes our approach to modeling stress paradigms suitable for neuroimaging environments. We showed a short film (4.54 minutes) with high negative emotional valence and high arousal content to 24 healthy male subjects (36.42 years old; SD=12.14) during fMRI. Independent component analysis (ICA) was used to identify networks based on spatial statistical independence. Through this analysis we identified the sensorimotor system and its influence on the dorsal attention and default-mode networks, which in turn have reciprocal activity and modulate networks described as emotional.

---

Understanding how the human brain processes information in a natural environment reveals information about its integral functioning during daily life. Under conditions simulating normal life) certain areas of the brain works as networks and participate in coding a complex stimulus, such as natural vision^1^; this brain activity reflects characteristics similar to those evoked in the natural human environment.

The vast majority of previous neuroimaging experiments sought to better understand the neural basis of perception using carefully controlled stimuli and tasks^1,^^2^. fMRI has been used to measure brain activity, primarily in the context, during highly controlled experiments using extremely carefully design stimuli^3^. In these cases, researchers use pre-determined, static, and isolated-object images that are flashed on the screen during purposefully predetermined paradigms that are synchronized to the image acquisition process^4^. Many contemporaneous experiments reduce the temporal complexity of their visual or auditory stimuli, presenting stimuli for one or two seconds or less. Yet, in order to sense and act in real-life circumstances, the brain must gather information over both long and short time intervals^5^. The use of precisely parameterized stimuli is critical for isolating the experimentally relevant dimensions from the extremely multidimensional natural stimuli.

Thus, in general, the world seen in the highly controlled fMRI experimental settings seems to bear little resemblance to our natural viewing experience ^4,6^. Because many studies use simplified static stimuli, surprisingly little is known about how the human brain operates during real-world experiences. Exploring brain function with dynamic, naturalistic stimuli is important for several reasons. First, it is vital to determine whether results obtained in experiments using simplified stimuli hold true under natural conditions. Second, some research questions can only be addressed with naturalistic tasks where there is little temporal regularity^7^. For these reasons, using more natural stimuli helps to detect brain activation patterns that are difficult to observe using simple stimuli and enables us to study the human brain under ecologically valid, naturalistic stimulus and task conditions^1^.

Through ICA connectivity approach it is proposed to explore the brain network implicated during an emotionally aversive condition. Studies about the stress impact on resting state networks (RSNs) show that stressed participants exhibited greater activation in the RSNs: Default Mode Network, Ventral Attention Network, Dorsal Attention Network, primary visual, and sensorimotor (SMN) contrary to non-stressed participants^8,9^. Also, participants in a stressful condition have evidenced deficiencies in the deactivation of RSNs vs. non-stressed participants^8^. Similarly brain connectivity during acute stress is related to alterations in brain areas related to perception, vigilance, and deactivation of the DMN, suggesting the promotion of focused attention that optimizes threat detection^10^. Literature shows that there are a few studies where ICA has been applied to observe the neural networks during emotion processing^11,12, 13,14^. Specifically there is a lack of studies that describe the active networks associated to resting state conditions under a stressful or emotionally disturbing condition^15,16^ considering an ecologically valid and naturalistic stimulus. Through this perspective it is proposed that the connectivity patterns evaluated reflects brain-environment interactions rather than stimulus responses^17^ as observed in classical experiments on stress processing. We used ICA approach as a valuable tool for revealing novel information about functional connectivity^18^ related with the stress processing phenomena.

The present research proposes studying the neuronal bases of the emotional processes in natural and free conditions without establishing an assumption or a controlled variable a priori. The aim is to validate an induced-stress method using neuroimaging paradigms. We propose a connectivity approach (independent component analysis-ICA) to characterize the brain networks implicated during the view of a stress-inducing video. In ICA, the fMRI activity is treated as a mixed signal that is mathematically divided into several, statistically separate signals known as independent components. These independent components can then be related back to the stimuli to understand the relationship between brain activity and stimuli^7^. So far ICA has been applied to fMRI data collected during movie watching^19,^ ^20^. In fMRI studies, ICA has been used to extract networks of brain activity during studies of baseline cognitive states and natural vision. Of the latter, those in the field of emotions use a connectivity approach that is based on the transient nature of the BOLD signal to estimate the connectivity networks^1^. The present study aims to identify the main connectivity networks and to elucidate their contrasts and the modulation that takes place during aversive emotional processing during a free-viewing paradigm. We approached this question by studying the functional organization of the human cortex when free-viewing a continuous sequence (4.54 minutes) taken from an original audio-visual feature film. We reasoned that such rich and complex visual stimuli are much closer to an ecological vision than the controlled stimuli usually used. Using a behavioral model and tests, we also evaluated whether presenting the video clip is a valid model for inducing psychological stress within an fMRI environment^21^. This video clip has been used in previous studies to induce a cognitive and emotional deficit^22,23,24^. However, the neural correlates linked to these psychological responses are unknown. Therefore, in the present study we also link the behavioral responses (perceived stress) derived from specific parts of said clip with increase connectivity in specific brain structures. To show that the results are specific to our particular experimental condition and do not merely reflect generic modulations in the audio-visual stimulation, a control experiment was carried out with a comparable set of “neutral” movie segments in an independent group of subjects.

## Results

### Behavioral data

The results show the statistically significant differences in the groups before (t1) and after seeing the video (t_2_). The scores (*M* = 2.77; SD = .59) at t2 are those with the highest stress subscale intensity (*t* = -6.99, *gl* =54, *p*<0.001) when compared with t1 scores (*M* = 2.09; SD = 0.56); this indicates that the stress-induction manipulation was successful.

No statistically significant differences at perceived stress were found in the group control before (t1; *M* = 1.74; *SD* = .58) and after seeing the movie’s segments (t2; *M* = 1.74; *SD* = .63) (*t* = .000, *gl* =30, *p* = 1.000) indicating a null influence of the control stimulus in the stress levels.

### fMRI ICA data

An estimated total of 31 components were observed by ICA (31 spatial maps with their corresponding time course) and ordered by the extent to which they explain the total variation in data. Five components were excluded (C17, C23, C24, C28 and C29) because they contained elements characteristic of a typical artifact^4^. The explained variance percentage value for the first component was around 6%, and it decays for later components, with the value for last component being 1.13%. On the other hand, in the average power spectrum of the time course of all the components, a predominant frequency was identified at 0.05 Hz, indicating the reliability of the components^25^. Computerized adjustment tests were performed in the FSL program MELODIC in order to determine how well the analysis of the group of components represents the joint activity of all the subjects. The results of these tests show the variety of relative responses in the whole session/subject analysis. The components have been ordered according to the mean response per decreasing component, without considering artifact components (17, 23, 24, 28 y 29), and all adjustments were significant (*p* < 0.00) for all except 30 (*p* < 0.00278) and 31 (*p* < 0.00268); the significance test was not corrected. Therefore, the trustworthiness of each component is known with regard to its representation of the joint activity.

### Network dynamics

A matrix correlation was built from time courses of every component (27) (Figure 1). The first 11 components had a significant correlation (*p* < 0.01) and were taken for further analysis. These represent diverse networks. C1, C2, C3 and C5 were identified as the same network (dorsal attention network), by having a similar hemodynamic pattern in their spatial maps, and having this high correlation. These four signals were averaged, obtaining a unique network for further analysis (Figure 2). Networks previously described as resting state^26^, as sensorimotor (SMN), default mode (DMN) and dorsal attention network (DAN) (Figures 2-3) were identified. The first 11 components of the matrix had a significant correlation (*p* < 0.01) and were used for further analysis. The networks represented by these components are shown in Figure 2.

**Figure 1.**
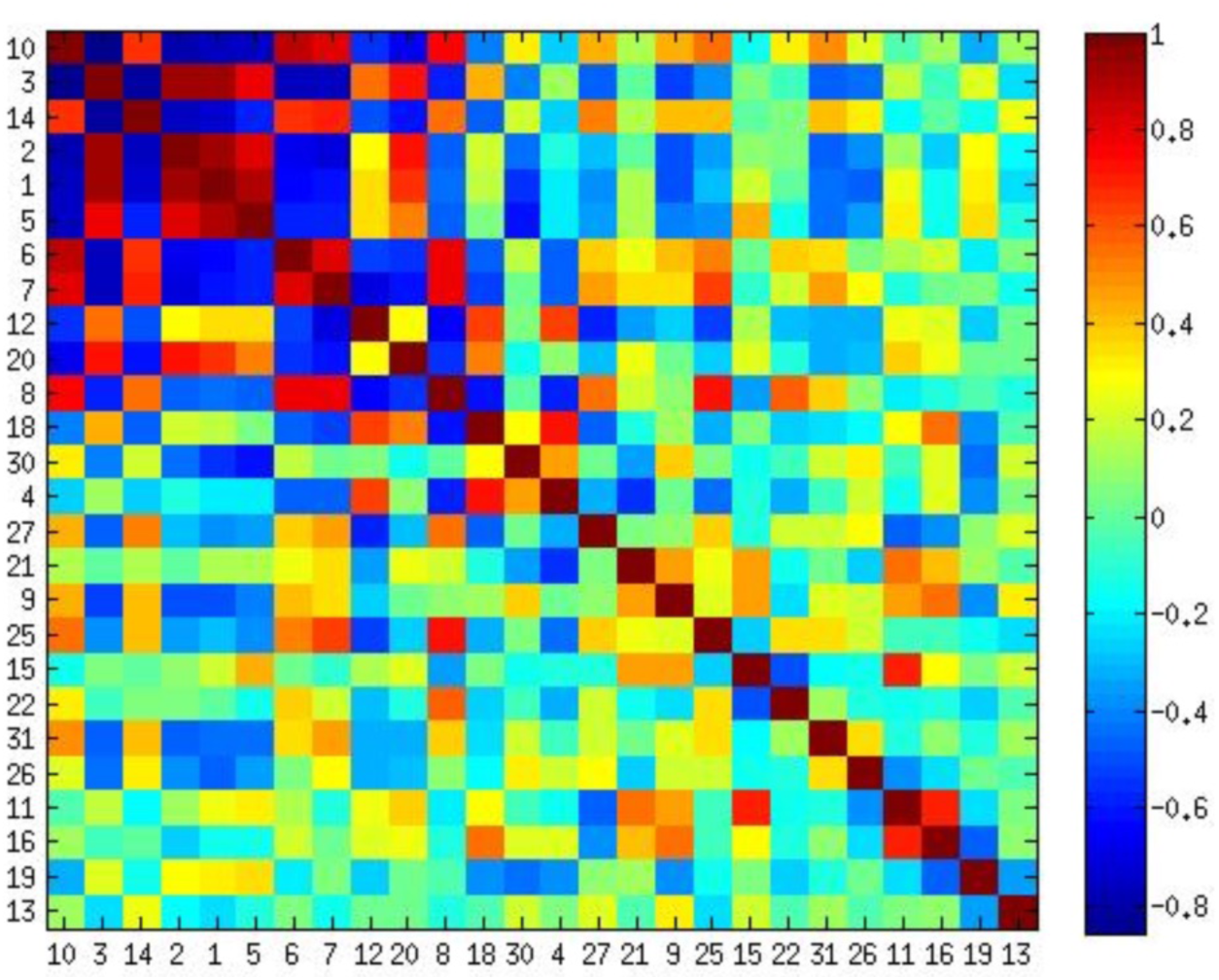
Correlation matrix of the BOLD signal time courses for the 27 ICA components. The matrix elements (I,j) represents the Pearson’s coefficient resulting from the cross correlation of the BOLD signal time course between the i and j components. the axes represent each of the component index, the color scale shows the Pearson’s value. Associated network activity is depicted in figure 2. Outline box contains the components which have a significant correlation (p<.01) taken for further analysis.

**Figure 2.**
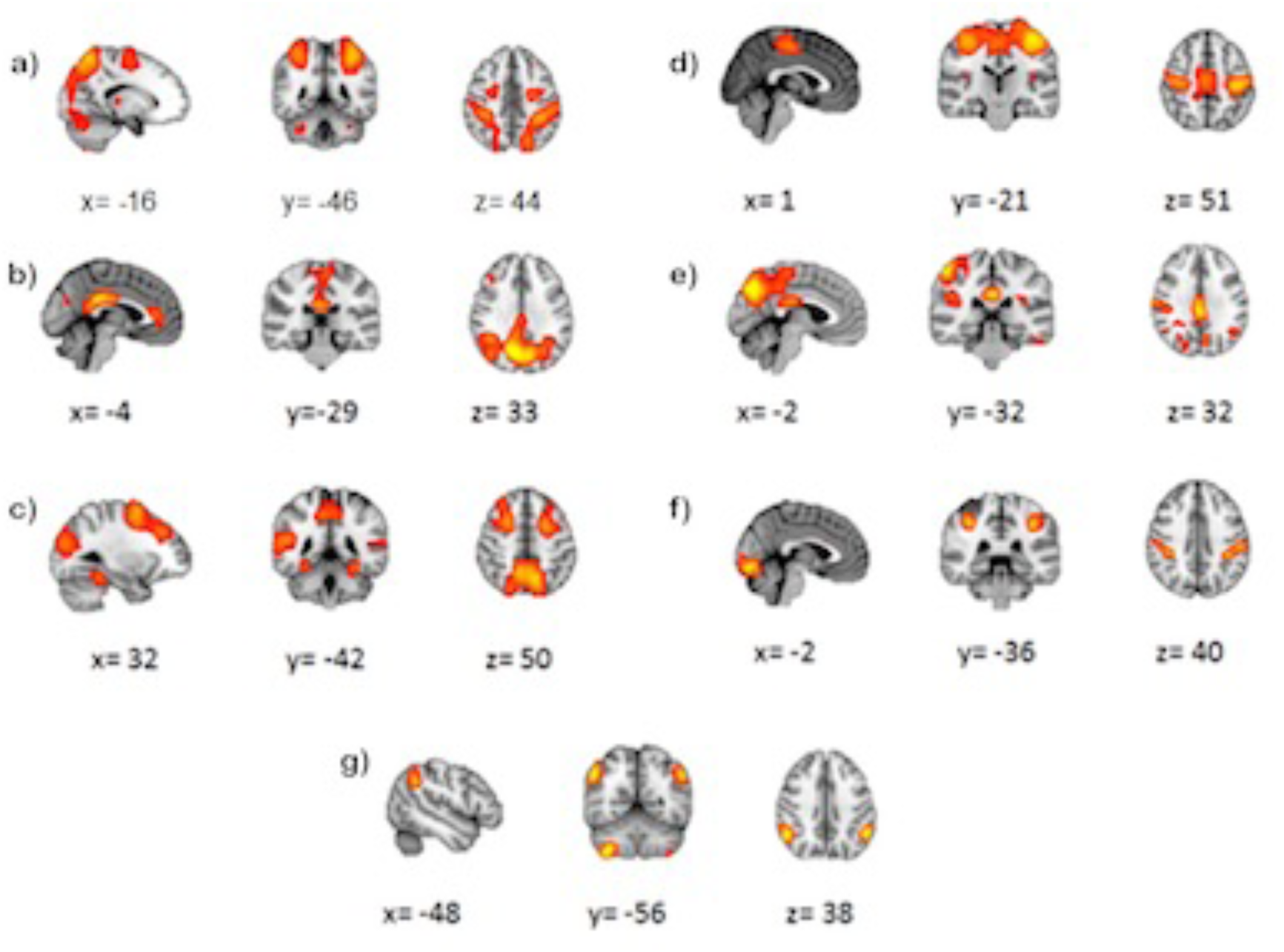
Some of the networks taken from the correlation matrix in figure 1 (outlined box) based on its order: a) C1, C2, C3 and C5; Dorsal Attention Network (DAN). b) C14; Sensory motor Network (SMN). c) C6; Default Mode Network (DMN). d) C7 posterior cingulate gyrus, lingual gyrus, precuneus. e) C12 Medial frontal gyrus, precuneus, medial temporal gyrus, fusiform gyrus. f) C20 Lingual gyrus, supramarginal gyrus, postcentral gyrus. g) C8 cerebellum anterior right lobe VIII-B area, angular bilateral gyrus.

**Figure 3.**
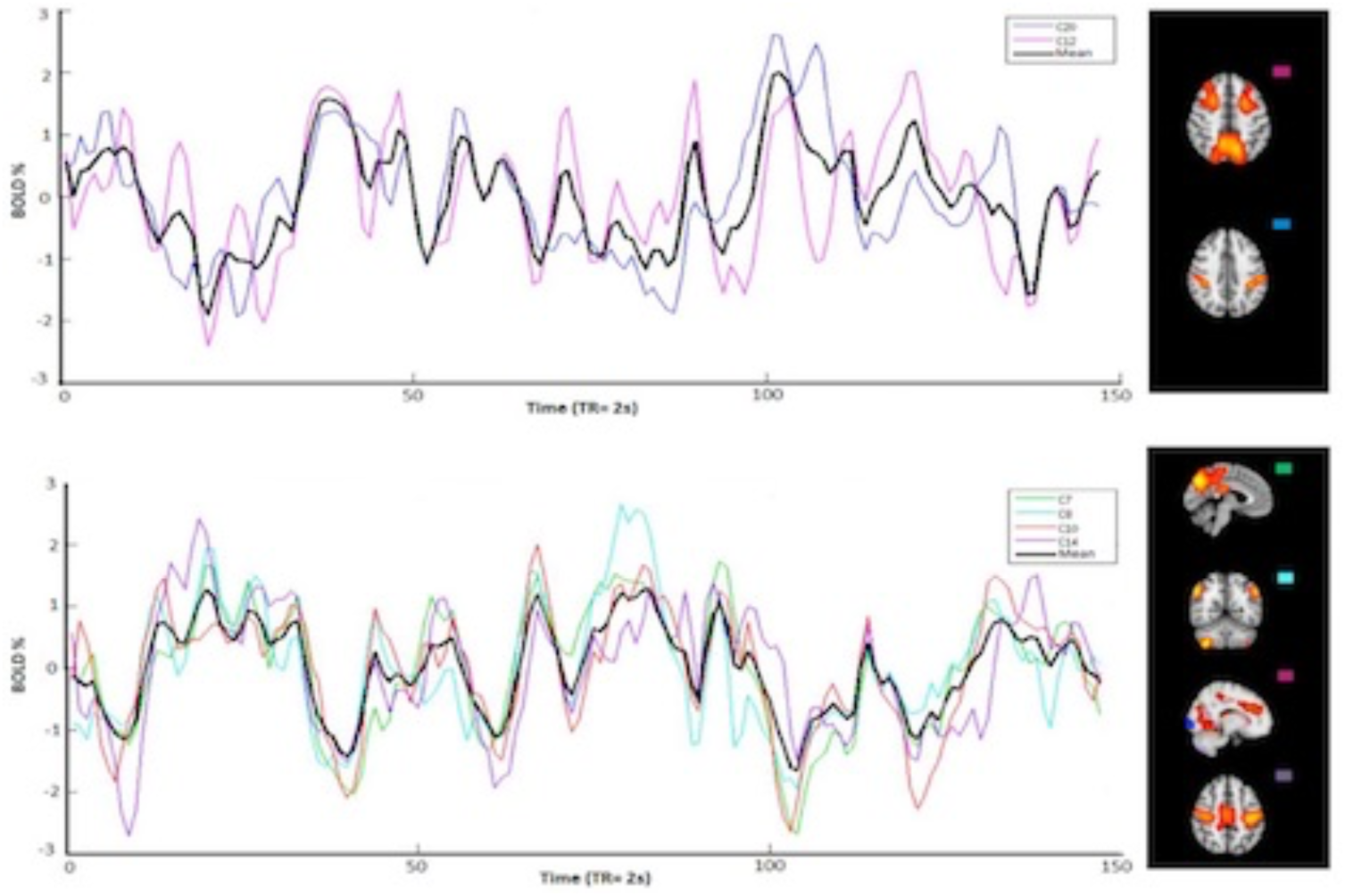
It shown components with significant correlation (see Fig 1). Average signals (black) obtained from BOLD signals. a) Components 12 and 20. b) Components 7, 8, 10 and 14.

We took the DMN and the DAN as time series references, since their activity has been better characterized in fMRI studies^27, 28, 29, 30, 31^. For demonstration purposes, two groups were formed based on their time courses. One group, comprising C7, C8, C10 and C14, has activity reciprocal to DMN. The other group, formed by C12 and C20, has activity reciprocal to DAN (Figure 3). The time courses of each group were averaged, resulting in two unique signals. By splicing these signals, a remarkable negative correlation between them was observed (*r* = -0.8452, *p*< 0.01) (Figure 4).

**Figure 4.**
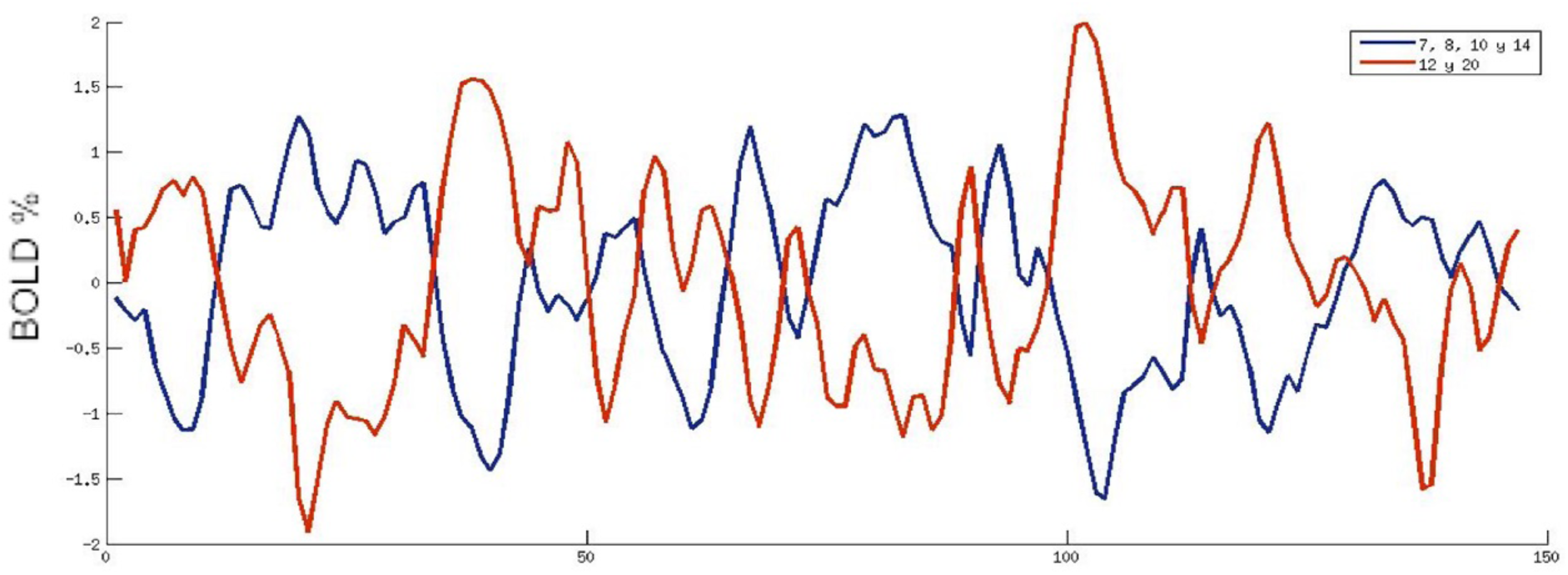
Two signal examples. The red line represents the average of components time courses 12 and 20 **group A**; the blue line of components 7, 8, 10 and 14, **group B**. The axes indicate the normalized BOLD signal (X) over time (Y).

As we know, changes in these time courses depend on the video display; thus, we ask what movie frames elicited the highest activation in these two signals. Figure 5 shows the movie frames from the highest activation peaks. The frames were ordered according to descending signal amplitude. Group a) was triggered mainly by scenes from the time prior to hurting the animal, whereas group b) was activated by scenes where the animal was dying.

**Figure 5.**
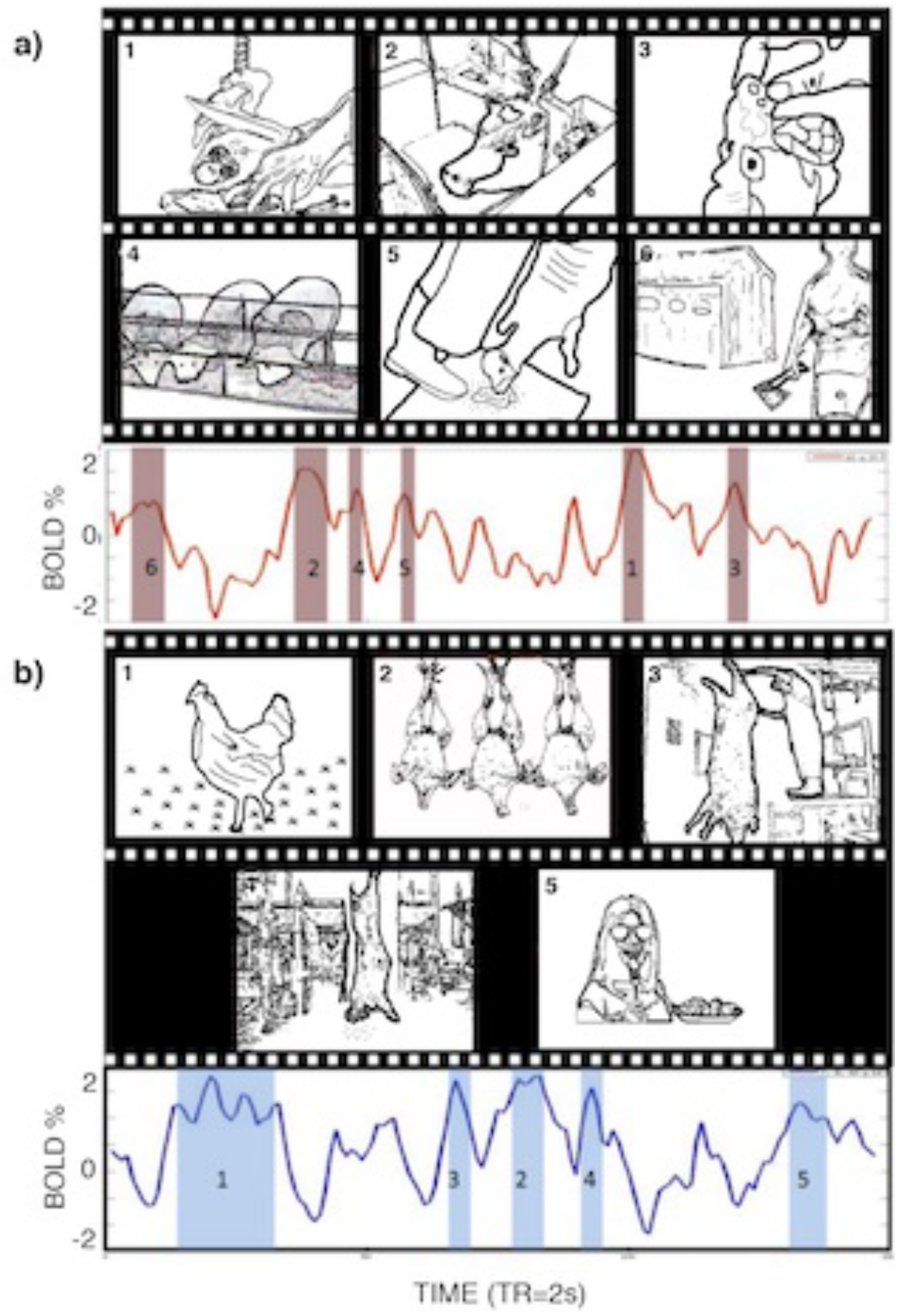
Types of scenes corresponding to high BOLD activity a) Scenes corresponding to the highest BOLD signal peaks, from group A (C12 and C20 signal), these images are related to the slaughtering actions and are images where the animals are prepared to be hurt. b) And from group B (C7, C8, C10 and C14 signal) while animal is hurting (in the dying process). Note that due to copyright reasons, the images showed in this figure are not the actual used in the study. However, all the images used in this figure represent similar situations observed in the original video. All the images used in this figure were constructed from images acquired and processed in our laboratory.

### Hierarchical modulation of networks

Our main objective was to elucidate the hierarchical modulation of networks between nodes (inter-modular) in a cognitive process. The results of the crossed correlation are shown in figure 6. Standing out among these findings is the SMN, which positively regulates the group of networks on the left side (DMN), and negatively regulates those in the right column (DAN). These two networks positively regulate other networks.

**Figure 6.**
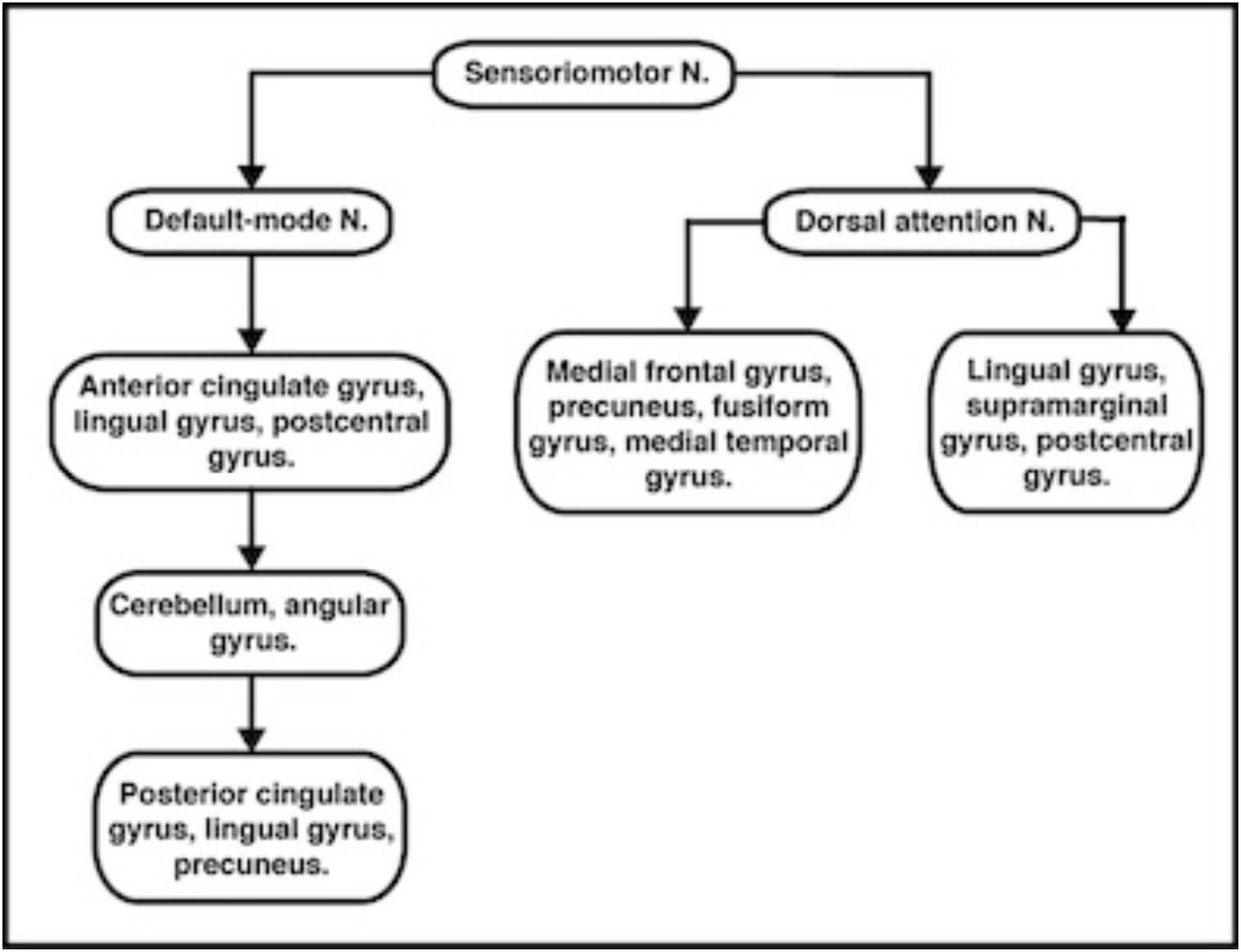
Networks organized hierarchically, based on the temporary delay and correlation that exist amongst them. The type of modulation is represented (positively and negative). Abbreviations: Sendoriomotor Network (SMN); Default mode network (DMN); Dorsal Attentional Network (DAN); Anterior Cinggulate Gyrus (ACG); Lingual Gyrus (LG); Postcentral Gyrus (PostG); Medial Frontal Gyrus (MFG); Precuneus (PCUN); Fusiform Gyrus (FUS); Medial Temporal Gyrus (MTG); Supramarginal Gyrus (SMG); Cerebellum (CER); Angular Gyrus (AG); Posterior Cingulate Gyrus (PCG).

Also observed were neural networks related to the induction of an emotional deficit, stress in this case; some of them were related to the limbic system or emotional processing areas such as a) posterior cingulate gyrus, lingual gyrus and precuneus; b) anterior cingulate cortex, lingual gyrus, postcentral gyrus and posterior insula; c) SMN, whose connectivity changed immediately after the animal-slaughtering scenes were shown. On the other hand, areas of networks with increased activity during scenes that predated the time of death, were a) medial frontal gyrus, precuneus, medial temporal gyrus and fusiform gyrus; b) lingual gyrus, supramarginal gyrus and postcentral gyrus. The analysis revealed that the high correlation is mostly present in posterior areas meanly the precuneous surface cortex, fusiform gyrus, follow by lateral occipital areas, lingual gyrus, as well as parts of dorsal limbic system (Fig. 7)

**Figure 7.**
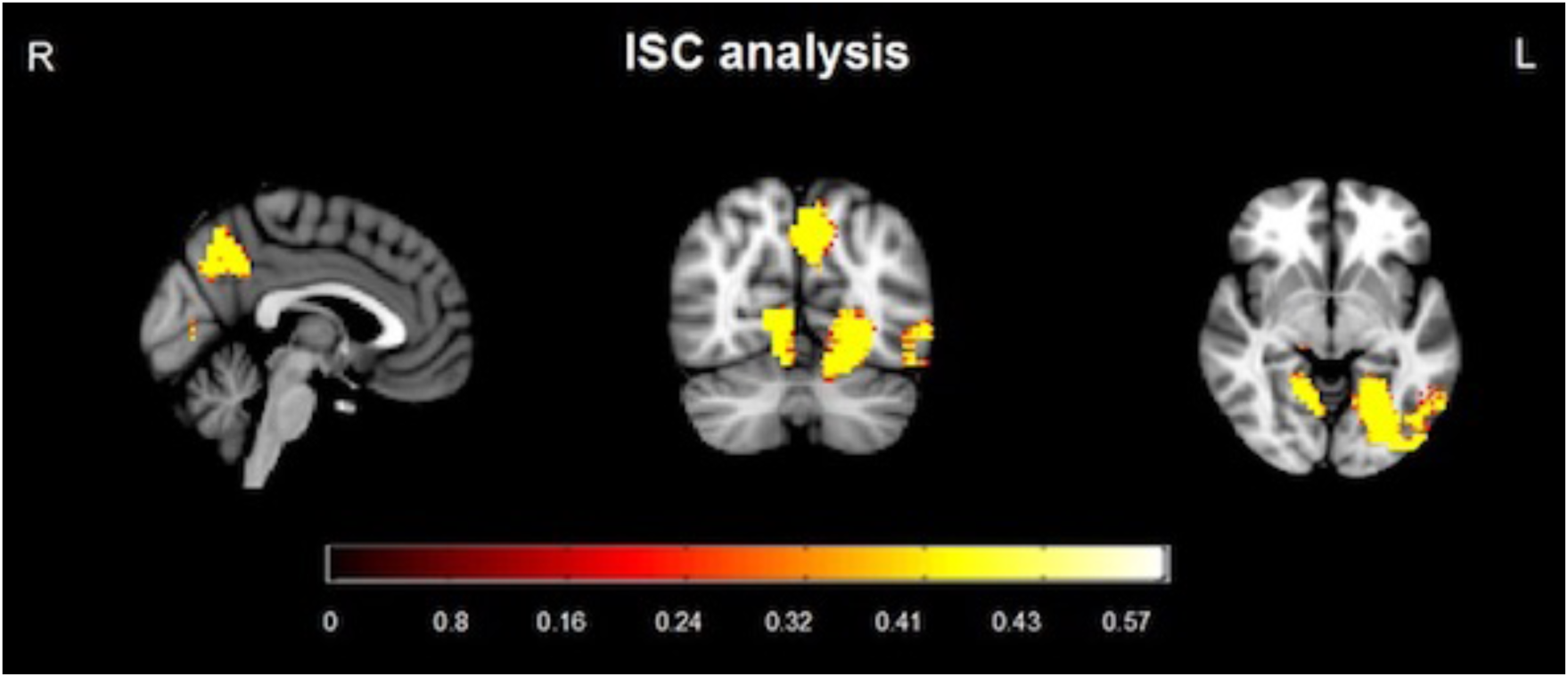
Shows the areas with higher correlation between subjects during the visualization of the video. Precuneous surface cortex, fusiform gyrus, lateral left occipital areas, lingual gyrus, and posterior cingulate cortex.

We can observe two clusters with the higher correlation, the precuneous cortex and the lingual gyrus. That means the activity in those areas it was present in a most synchronized manner in all subjects across the time. If we compare with ICA results, the lingual gyrus appears in three of the networks (components 7, 10 and 20) related with emotional scenes as it could observe. Precuneous is part of the component 7, also an important part forming the default mode network, together with the posterior cingulate cortex, combines bottom-up attention with information from memory and perception.

## Discussion

The present study was designed to locate the main networks, their dynamics, and the modulation that takes place in aversive emotional processing during a free-vision model in order to validate a stress-inducing method within a functional magnetic resonance environment. Using ICA we were able to document the dynamics that exist between the BOLD activity of each network detected while a stressful video was being viewed. What stood out the most in this study was the identification of the DAN and the DMN. Our findings are consistent with previous research, especially the determination that these networks are intrinsically anti-correlated^32, 33^. Furthermore, the signal of each network is also associated with the reciprocal activity of other networks of the six networks related to emotional processing. Based on the correlation of their time courses, these six networks formed two subgroups, where the average signal of each subgroup had a high negative correlation value (r= -0.8452). The first subgroup, made up of components 12 and 20, had networks that involve regions, such as the medial frontal gyrus, precuneus, middle temporal gyrus, fusiform gyrus, lingual gyrus, supramarginal gyrus, and postcentral gyrus, that have reciprocal activity with the DAN and are observed in moments of preparation before killing an animal. This coincides with processes associated with this specific network, which are expectation and fixed attention when knowing that something is going to happen soon, marking a striking difference from other processes related to attention^34^. During this video, the subjects know the nature of the video, but they do not know for certain what will happen or how. So, their attention is sustained during this whole process, linked to an emotional network that contains the supramarginal gyrus and the postcentral gyrus, which are associated with emotional states to others, including pain and suffering. This signal change happens even without the stimuli that provoke an emotional response; it is triggered by knowing that they will happen. This activity begins to decline the moment the expected event occurs. From this point on, activity begins for the following group of networks. Prior findings are consistent with the responses related to emotional stress obtained in behavioral registers, validating the efficiency of this method to induce stress. The ICA components 7, 8, 10, and 14 include regions 7 (posterior cingulate, lingual gyrus, and precuneus), 8 (right frontal lobe of the cerebellum area III-B and bilateral angular gyros), 10 (anterior cingulate, lingual gyrus, post-central gyrus and posterior insula), and 14 (sensorimotor system, pre- and post-central gyrus). The corresponding BOLD activity involves the DMN. Despite the fact that the DMN is characterized as being present mostly during a resting state, its activity also persists during states of sensory processing that impose a minimum cognitive demand^35, 36^. An example of this is a video where the subject does not have to perform any action. Additional aspects and functions of the DMN have been coming to light that underlie this network’s activity, such as: its alteration in different disorders (epilepsy, schizophrenia, depression, substance abuse, etc.), or the modulation of emotional processes, as demonstrated here. A modular focus of the brain tells us about its capacity to assemble quickly and robustly, in order to segregate an infinity of processes. This idea has been developed more tangibly using computer mechanisms that demonstrate its high efficiency and functioning by modulating multiple processes of complex information and developing tasks in a changing environment^37^. The main and most important networks for the modular control of the brain have been demonstrated before and include: the occipital, sensorimotor, and default mode networks^38, 39^. These same networks were found at the hierarchical level of modulation^40^, whereas at the voxel level researchers found visual, auditory, and default mode networks and subcortical areas^41^.

Additionally, this study showed the importance of the SMN and DMN. As was made visible in the results of the crossed correlation, the sensorimotor network precedes both of these groups in activity, and its activation in the absence of pain or physical sensation is associated with the perception of others’ pain. This may indicate that perception of other’s pain has greater significance than other brain structures activity, due to the high negative valence and high arousal of the stimulus presented; thus, the network responding to perception of other’s pain should be the main modulating center.

The moment the injury scene starts, the sensorimotor network begins to activate and the attention network begins to deactivate, which indicates negative modulation. Hence, tension control is directed by the stimulus. Afterwards, this same network (sensorimotor) gives way to the DMN, which modulates other emotional networks areas such as the lingual gyrus, which has numerous connections with the amygdale, central lobe, (associated with the perception of unpleasant stimuli), the anterior and posterior cingulated cortex, part of the limbic system, cerebellum, postcentral gyrus, and angular gyrus.

Thus, we may summarize our data by analogy to a basic brain process. The brain process always has relay points, starting from the first, the most basic, up to however many relays the complexity of the process may require. In this case we may speak of relay networks instead of relay areas, where the basic network (sensorimotor) is always activated by these stimuli, and, depending on the type of stimulus, it may activate the default mode or the attentional network, which then continues with networks that process emotions and so on, depending on the complexity of the stimulus, just as any other brain process. The present study also showed that the same modulating networks are maintained in both a stable state and a stimulus-triggered altered state, leading to the possibility that these networks act as part of the general modulation of brain activity, regardless of the stimulus or process. However, this conclusion must be confirmed by studies made under the same model with different types of stimuli that provoke responses related to other cognitive processes. The present findings suggest the usefulness of the method for inducing stress during neuroimaging studies. We have explored the human brain’s response under stress considering a dynamic naturalistic stimulus^5^. Our technical and methodological approach differs from previous studies of psychological and physical stress paradigms suitable for neuroimaging environments^42, 43^. Further, using more natural stimuli will probably allow us to detect patterns of brain activation that are difficult to observe using simple stimuli, and enable us to study a “stressed human brain” under ecologically valid naturalistic stimulus and task conditions^1^. This approach is a cost-effective and efficient way to elucidate the mechanisms underlying stress. Our study investigates the neural circuitry of stress in combination with psychological stress measures. Future studies would consider additional measures (physiological) of stress together with this method.

## Methods

### Participants

Twenty-four right-handed healthy male subjects were enrolled in the study. All subjects gave their written informed consent, the experimental protocol was approved by the Bioethics Review Board of the Institute of Neurobiology, UNAM, and was performed in accordance with the Declaration of Helsinki. Subjects had an average age of 36.42 (*SD* 12.14) and a minimum of 12 years of schooling. All subjects were mid-to-high socioeconomic level. Individuals who met any of the following criteria were excluded: history of head injury, treatment with psychotropic medications, use of narcotics, steroids, or any other medication that affects the central nervous or endocrine systems, having had a medical illness within 3 weeks prior to testing, self-reported mental (SCL R-90; ^44^) or substance use disorder, daily tobacco use, regular nightshift work, current stressful episode or major life event, and regularly viewing extremely violent movies or playing violent computer games. One subject was excluded during the analysis due to numerous motion artifacts. Women were excluded from this study since the hormonal cycle may influence emotional responses and may cause a bias in the study^45^.

### Stimuli and procedure

Prior to entering the scanner, subjects received thorough instructions about the scanning procedure and the tasks to perform. Perceived stress (aversive stimuli) was tested immediately after entering the scanner to obtain a baseline level (t_1_) and again after viewing 4.54 minutes of the stressful movie (t_2_).

### Psychological measures

Perceived stress. The computerized version of the Stress and Activation Adjectives Checklist-SAACH^46^ was employed, considering only the stress dimension (11 adjectives with a four point answer format: absolutely true, probably true, not sure and absolutely not). In the present study the scale was adapted to be administered in the scanner environment. To respond to the test, subjects had a magnetic resonance (MR) compatible ResponseGrip (NordicNeuroLab) with their left and right index and thumb fingers each placed on one of four buttons. All sentences and instructions were presented on a GUI programmed in Java environment running on a MS Windows platform. Participants were instructed to rate their present emotional state. The items presented were randomized. The sentences were printed in lower case letters, displayed in approximately 18-mm tall Arial, white on a black background, providing high legibility. Prior to the scanning session, each subject was instructed in the performance of SAACH and the use of the computer mouse by practicing a shortened version outside of the scanning environment on a desktop computer with verbal instruction from the researcher.

### Aversive stimuli

We use fMRI coupled with the eye tracking technology to confirm attentive viewing of all projected stimulus and movie fragments. The eye tracker was calibrated for each participant before the experiments began. A MR-compatible NNL eye-tracking camera was used to record video data of the subject’s eyes during the fMRI task-related scanning (Visual System NNL EyeTracking Camera, and ViewPoint EyeTracker software, Arrington Research Inc., Scottsdale AZ). Subjects watched 4 min and 54 sec of the English movie “Faces of Death #1” (dir. John Alan Schwartz, 1978, original length 105 min) in the fMRI scanner. Three short movie fragments were used to create the proper context (1 × 60s, 1 × 120s, 1 × 114s). This shorter version was re-edited (co-author, LG^*^) with computer software. Selected fragments were comparable in amount of speech, human presence, luminance, and language. The aversive movie clips contained scenes of a woman on a farm decapitating a rooster, images of a slaughterhouse where sheep and cattle are being sacrificed in a brutal manner and video fragments of a group of actors pretending to eat the brain of a monkey that they had to kill themselves. Participants were instructed to carefully watch the video and feel free to stop the video if the images were disturbing them. Participants were informed before the experiment that watching the film might be stressful and that they could terminate the experiment at any point. The video clip image contents were classified with negative valence and high arousal content according to Bradley y Lang^47^.

### Functional Magnetic Resonance Imaging

During projection of the video, the sequence of images was captured with a 3.0 Tesla Discovery MR750 MRI unit, using the 32-channel reel for cranium, at the Neurobiology Institute of the UNAM. The functional images were acquired with a sequence of Eco Planar (EPI) pulses for heavy images at T2^*^, GE-EPI of TR / TE = 2000 / 40 ms in a 64 × 64 matrix over a FOV of 25.6 cm in 36 slices with a width of 4 mm per slice. This resulted in isometric voxels with a spatial resolution of 4 × 4 × 4 mm^3^. The high-resolution structural images were acquired using a T1 weighted SPGR pulse sequence with 1 × 1 × 1 mm^3^ of spatial resolution.

## Data Analysis

### fMRI Data

All data were transferred to offline work stations to convert them from DICOM to NIfTI format (dcim2nii, Ch. Rorden, <u>http://www.nitrc.org/projects/dcm2nii);</u> afterwards they were processed using the MELODIC ICA module of the FSL program, which runs an algorithm of independent components per voxel in the stack of images for the whole group of subjects. It was used MCFLIRT tool for motion correction^48^, timing correction for interleaved slice acquisition, spatial smoothing of 5mm and low-pass filtering. For a fit to standard space: MNI 152 Talairach^49^ linear normal search with 12 degrees of freedom, we used 0mm of resampling resolution for single-session ICA and 4mm for Multi-session Tensor ICA. It was applying variance normalize time-courses, automatic dimensionality estimation, threshold IC maps: 0.5, and mean high resolution for background image. FastICA algorithm was used as well the spatial ICA method approach.

The first analysis was made using the *Single-session ICA* mode that makes up a standard of each entrance file (23) to a two-dimensional matrix (time and space) in order to decompose them later into two-dimensional matrixes (time and components, space and components). The result estimates the independent components for each subject without taking others into account. The results of the “single session” were used to identify the components that corresponded to each subject’s noise, and these were eliminated (between four and five per subject), thus arriving at a more reliable database^25^. For the second level of analysis in MELODIC, it was used “Multi-session Tensor-ICA” which takes the entry data as a three-dimensional matrix (time, space, and subjects) and breaks it down into triplet, two-dimensional matrixes: time courses-ICA components and spatial maps-ICA components. The final observable product describes common components in all or most of the subjects and orders them according to the highest percentage of variation explained by the model, optimized by a distribution of non-Gaussian spatial sources, using fixed point iteration^50^. The resulting components went through Pearson correlation tests and linear regressions were calculated, to determine the network dynamics and the modulation between networks^51^, linear regression was used to measure the strength and direction of the linear relationship between two joint signals^52^. All these analyses were processed using our own written programs in MATLAB (Mathworks, Natick, MA, USA).

To support ICA analysis, and since GLM is not suitable in this case, we used Inter-subject correlation analysis, which is highly comparable with GLM^53^. ISC analysis allows visualizing selective and time-locked activity across a wide network of brain areas, under some natural stimuli, comparing the whole neural response across all subjects, without a model^54^. We used the same data of the 23 subjects, normalized into an MNI coordinate system, spatially smoothed and realigned. The ISC analysis was performed using ISCtoolbox for Matlab by Kauppi et al^55^ (<u>http://code.google.com/p/isc-toolbox/).</u>

### Behavioral data

To test whether the video caused aversive psychological effects, behavioral data was analyzed with paired sample *t*-tests comparing stress measures between t_1_ and t_2_.

## Neutral movie condition

### Participants

The participants were 31 students (21 women, 10 men; *M* age = 20.90 yr., *SD* = 5.32) from the Psychology Department of Universidad de Guanajuato, who gave their informed consent to being part of the study. One participant lacking complete data was omitted (final n = 32).

### Psychological measures

Perceived stress was measured with the pencil-paper version of the Stress and Activation Adjectives Checklist.

### Neutral stimuli

The subjects watched 17 neutral fragments of the same movie projected in the previous study with 4 min and 53 sec length (1-17 x 16 s). This neutral version of the movie was reedited and implemented thorough Java software. The neutral movie clips contained colored emotionally neutral scenes (e.g. hands, one house, sheeps and cows in its ordinary context and scenes of people in abstract or trivial circumstances).

### Procedure

Perceived stress was tested before (t1) and after viewing (t2) the neutral fragments. All video stimuli were presented via a video projection system in a semi-darkened experimental room isolated from noise and distractions. This system projected a 1.2 x 1.8 image onto a reflexive white screen. For all experimental stimuli, the picture took up 80% of the screen on a black background. The stimuli were evaluated in one single experimental session and 15 minutes of length.

### Data Analysis

Behavioral data was analyzed with paired sample *t*-tests comparing stress measures between t1 and t2.

## Acknowledgements

We thank the Programa de Maestría en Ciencias (Neurobiología) of the Universidad Nacional Autónoma de México (UNAM) for the masters fellowship (MN) supported by the National Council of Science and Technology, Mexico (Consejo Nacional de Ciencia y Tecnología, CONACYT); DGAPA-UNAM for the Posdoc Fellowship (JM-S) the PASPA program at DGAPA-UNAM for the sabbatical fellowship (FAB), and CONACyT for the funding received via grant CB167271 (FAB). We are also grateful to Dr. Erick H. Pasaye for technical support and Drs. D. Pless and M.C. Jeziorski for their revision of the manuscript.

## Author Contributions

JM-S and FAB designed the experiment, MN, JM-S, LG-S and FAB acquired all the data, MN analyzed the data, MN and FAB wrote the main manuscript text, and MN and LG-S prepared figures 1-6. All authors reviewed the manuscript.

